# ‘Munger’ black raspberry: Insights into its role as a virus indicator for Rubus species

**DOI:** 10.1101/2025.01.14.633050

**Authors:** Andrea Sierra-Mejia, Ioannis E. Tzanetakis

## Abstract

‘Munger’ black raspberry (*Rubus occidentalis*) has been a preferred indicator for *Rubus* viruses. The working hypothesis behind the ‘Munger’ elevated susceptibility to viruses is its ability to sustain elevated virus titers; however, no research has yet explored this concept. To address this, we utilized an infectious clone of blackberry chlorotic ringspot virus to study the differences in virus expression between two *Rubus* species: *R. occidentalis* (‘Munger’) and *R. L. subgenus Rubus* Watson (blackberry, ‘Natchez’). Our data demonstrate that virus accumulation in ‘Munger’ is over 47,300 times higher compared to ‘Natchez,’ enhancing our understanding of virus-host dynamics and providing valuable insights into why ‘Munger’ is a preferred indicator species for virus detection in *Rubus* spp.

## 1. INTRODUCTION

Symptom expression serves valuable diagnostic purposes. For this reason, scientists have identified susceptible hosts (indicators) that exhibit distinct symptomology when infected by an array of pathogens. A notable example is ‘Munger’ black raspberry (*Rubus occidentalis*), which was released in the late 19^th^ century (Hedrick, 1925) and is considered the best indicator for the majority of *Rubus* viruses (Martin et al., 2016; Martin & Tzanetakis, 2008) even though many of the ones discovered in the 21^st^ century tend to be asymptomatic (Martin et al., 2013; Villamor et al., 2022). Still, the factors contributing to its greater vulnerability to viruses have not been studied.

The research presented within was initiated based on a simple inquiry: Why has ‘Munger’ been the preferred indicator for Rubus viruses for over a century? To explore this question, we formulated a straightforward hypothesis: ‘Munger’ can sustain high virus titers over other *Rubus* species and cultivars.

To test this hypothesis, we required two essential components: virus-free plant material and a pure, homogeneous virus inoculum. This inoculum needed to be free from the inherent quasi-species diversity, titer variability, and potentially varying biological properties typically associated with grafting or mechanical inoculations of filed material. We obtained ‘Munger’ and ‘Natchez’ (*R*.*L. subgenus Rubus* Watson, blackberry) plants free of viruses based on HTS analysis (Druciarek et al., 2024). We selected the infectious clone of blackberry chlorotic ringspot virus (BCRV), a member of the genus *Ilarvirus* (Sierra-Mejia et al., 2025), as the model virus to compare virus expression between species over a six-month time period.

It will appear contradictory to use BCRV in the study given its asymptomatic nature in single infections across both host species (Poudel et al., 2011). The choice was deliberate and is grounded in the study’s hypothesis – ‘Munger’ sustain high virus titers compared to other *Rubus* species and cultivars. This approach also eliminates the confounding influence of symptoms, which could inherently skew virus titer comparisons between tissues (Quito-Avila and Martin, 2012; Ammara et al., 2017; Bin et al., 2022). The results of the study will help us deepen our understanding of virus-host interactions, essential for enhancing agricultural practices and improving plant health management.

## 2. MATERIALS AND METHODS

### 2.1 Plant maintenance

The plants selected for this study were confirmed to be singly infected with BCRV (Druciarek et al., 2024, Sierra-Mejia et al., 2025). BCRV-infected ‘Natchez’ plants were acquired using dodder to transfer the virus from *Nicotiana benthamiana* (Sierra-Mejia, 2024), whereas the BCRV-infected ‘Munger’ plants were obtained via direct agroinoculation. Infected plants were vegetatively propagated to increase their numbers. To establish a consistent baseline and minimize the impact of environmental variations and virus delivery, all plants were trimmed and subjected to chilling conditions by placing them in a cold room at 4°C in complete darkness for one month. The surviving plants—17 ‘Natchez’ and 8 ‘Munger’—were transferred to a growth chamber, where they were maintained under a 16/8-hour light/dark cycle at 22°C.

### 2.2 Detection

Sampling was conducted monthly over a six-month period. To minimize titer variation related to tissue age, we consistently sampled only newly expanded leaves. Total nucleic acid extractions and cDNA synthesis were performed as described by Poudel et al (2013). cDNA was diluted at a ratio of 1:4 and 1:4,000 for virus and 18S rRNA detection, respectively. All RT-qPCRs were conducted using a CFX96 Touch real-time PCR detection system (Bio-Rad, Hercules, CA, USA). For virus detection the thermocycler conditions consisted of 50° C for 2 min, 95° C for 10 min, followed by 40 cycles of 95° C for 15 sec and 60° C for 1 min using the primers/probe of Poudel et al (2014) (Supplementary Table 1). For 18S rRNA detection was performed using primers and probes designed by Osman et al (2007). The probe’s quencher and fluorophore were further modified to meet the specific requirements of the equipment (Supplementary Table 1). The final reaction volume was 20 µL and included 2 µL of cDNA, 10 µL of TaqMan™ Universal PCR Master Mix II with UNG (Thermo Fisher Scientific), 25 nM of each 18S rDNA-specific primer, and 12.5 nM of the corresponding probe. The thermocycler conditions were the same as those used for BCRV detection, and appropriate controls were included in each run. Each sample was analyzed in triplicate through qPCR, and the average CT value from the three independent runs was calculated (Table 1).

**Table 1.** RT-qPCR results for detecting blackberry chlorotic ringspot virus in ‘Munger’ and ‘Natchez’ plants over a six-month period. A given value indicates that the virus was detected, the number corresponds to the average CT amongst three independent runs, (-) indicates that the virus was not detected, a CT value of 40 was given to allow for the statistical analyses, and (NA) signifies that the plant was lost and could not be sampled for further analysis.

|  | Month |  |  |  |  |  |
| --- | --- | --- | --- | --- | --- | --- |
| Plant | 1 | 2 | 3 | 4 | 5 | 6 |
| **Ct Values- Munger** |  |  |  |  |  |  |
| **MD1** | 21.86 | 21.38 | 23.90 | 22.72 | 21.14 | 24.67 |
| **MD2** | 21.85 | 18.41 | NA | NA | NA | NA |
| **MD3** | 20.38 | 20.28 | 21.67 | 23.41 | 20.57 | 22.92 |
| **MD4** | 21.38 | 18.07 | 22.86 | 19.82 | 21.74 | 23.10 |
| **MD5** | 19.39 | 17.16 | 19.65 | 20.72 | 20.43 | 23.14 |
| **MD6** | 24.32 | 20.81 | 21.00 | 20.41 | 22.29 | 24.85 |
| **MD7** | 22.79 | 19.93 | 20.59 | 22.79 | 20.71 | 24.13 |
| **MD8** | 22.45 | 21.95 | 21.39 | 21.47 | 21.06 | 23.88 |
| **Ct Values- Natchez** |  |  |  |  |  |  |
| **ND1** | - (40.00) | 38.37 | - (40.00) | - (40.00) | - (40.00) | - (40.00) |
| **ND2** | - (40.00) | - (40.00) | - (40.00) | 39.25 | - (40.00) | - (40.00) |
| **ND3** | 34.53 | - (40.00) | - (40.00) | - (40.00) | - (40.00) | - (40.00) |
| **ND4** | - (40.00) | - (40.00) | - (40.00) | - (40.00) | - (40.00) | - (40.00) |
| **ND5** | - (40.00) | 38.73 | - (40.00) | - (40.00) | - (40.00) | 39.13 |
| **ND6** | - (40.00) | - (40.00) | - (40.00) | 38.90 | - (40.00) | 38.77 |
| **ND7** | - (40.00) | - (40.00) | - (40.00) | - (40.00) | - (40.00) | 39.48 |
| **ND8** | - (40.00) | 38.97 | - (40.00) | - (40.00) | - (40.00) | 38.91 |
| **ND9** | 36.12 | 39.68 | - (40.00) | - (40.00) | - (40.00) | - (40.00) |
| **ND10** | - (40.00) | - (40.00) | - (40.00) | - (40.00) | - (40.00) | - (40.00) |
| **ND11** | - (40.00) | - (40.00) | - (40.00) | - (40.00) | - (40.00) | - (40.00) |
| **ND12** | - (40.00) | - (40.00) | - (40.00) | - (40.00) | - (40.00) | - (40.00) |
| **ND13** | 38.52 | 37.55 | - (40.00) | - (40.00) | - (40.00) | - (40.00) |
| **ND14** | 33.14 | - (40.00) | - (40.00) | - (40.00) | - (40.00) | - (40.00) |
| **ND15** | - (40.00) | 37.27 | - (40.00) | - (40.00) | - (40.00) | 39.45 |
| **ND16** | - (40.00) | - (40.00) | - (40.00) | - (40.00) | - (40.00) | 39.23 |
| **ND17** | - (40.00) | 36.17 | - (40.00) | - (40.00) | - (40.00) | - (40.00) |

### 2.3 Statistical analysis

The resulting qPCR runs were compiled into a dataset, categorizing the data by month and species. Cycle threshold (CT) values were normalized for viral quantification using 18S rRNA as an internal control, and we included this value in our dataset. For normalization purposes, a CT value of 40 was assigned when the virus was not detectable. Normalization was calculated using the equation:

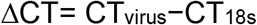

Where, ΔCT represents the difference in CT values between virus and internal control (18S rRNA). The normalized value, reflecting relative virus expression, was derived from:

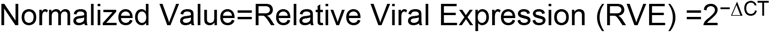

The dataset was imported into R version 4.4.1 for analysis (R Core Team, 2023) which was used in all downstream analyses. We employed a linear mixed effects model, with the normalized values as the response variable. Species and month were included as fixed main effects, and an interaction effect between month and species was also included. Each individual plant was accounted for as a random factor to address the variability amongst plants. A Type III analysis of variance (ANOVA) using Satterthwaite’s method was conducted to determine the significance of the fixed and interaction effects in a linear mixed-effects model. Following this, a post hoc analysis was conducted to address specific group differences in the mixed effects model. For this, marginal means were evaluated, and pairwise comparisons among these means were conducted using Tukey’s Honest Significant Difference (HSD) method (P < 0.05).

## 3. RESULTS

BCRV was consistently detected in all ‘Munger’ plants at all sampling points. In ‘Natchez’ the virus was undetectable at certain sampling points in some plants, without any consistent patterns emerging (Table 1).

Overall, the RVE over the length of the study was 0.109 for ‘Munger’ and 2.31e^-6^ for ‘Natchez’, yielding an RVE ratio of over 47,304. The ANOVA test showed that both species (p=7.47e^-10^) and sampling time (p=8.66e^-10^) had a significant effect on virus RNA accumulation (Table 2). Additionally, the interaction between the two was also significant (p = 8.64e^-10^), suggesting that the effect of species on the virus accumulation may vary depending on the month (Table 2). The post hoc analyses revealed significant differences in the estimated marginal means (emmeans) of virus accumulation between ‘Munger’ and ‘Natchez’ over the length of the study. For example, in the first month, the emmean for ‘Munger’ was 0.05, whereas for ‘Natchez’, it was 6.10e^-7^, indicating higher virus accumulation in the former. This trend continued in subsequent months, with ‘Munger’ consistently displaying higher emmean values (Figure 1). Furthermore, these analyses revealed significant differences in virus accumulation associated with the sampling time. For ‘Munger’, there was considerable variability, with a notable increase in virus expression during the second sampling time, whereas ‘Natchez’ exhibited less variability across sampling times.

**Table 2.**
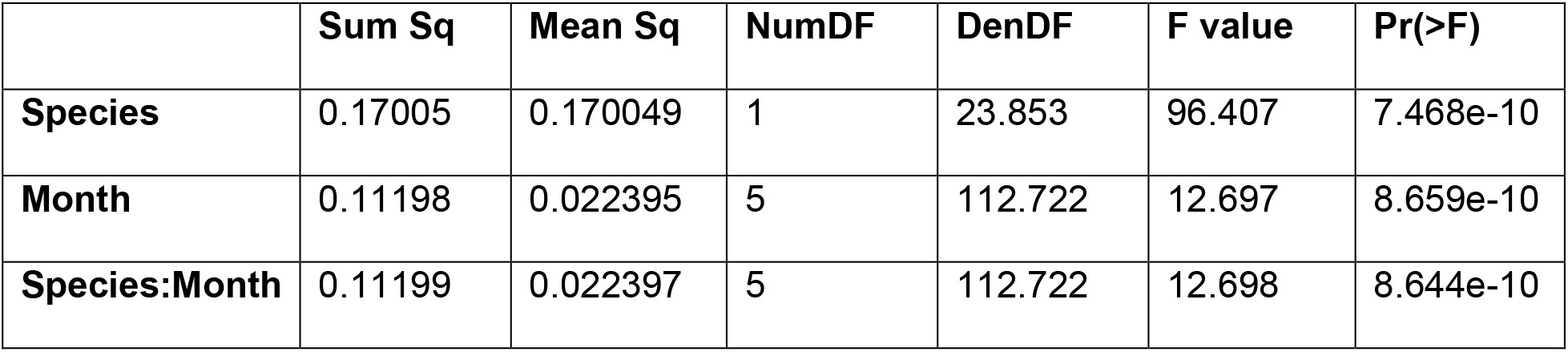
Type III Analysis of Variance with Satterthwaite’s Method. The table shows the results for the effects of species, month, and their interaction on the response variable. All factors (species, month, and species:month interaction) are statistically significant, with p-values less than 0.001. Sum Sq(Sum of Squares), Mean Sq(Mean Square), NumDF (Numerator Degrees of Freedom), DenDF(Denominator Degrees of Freedom),F value, and Pr(>F) (p-value).

**Figure 1.**
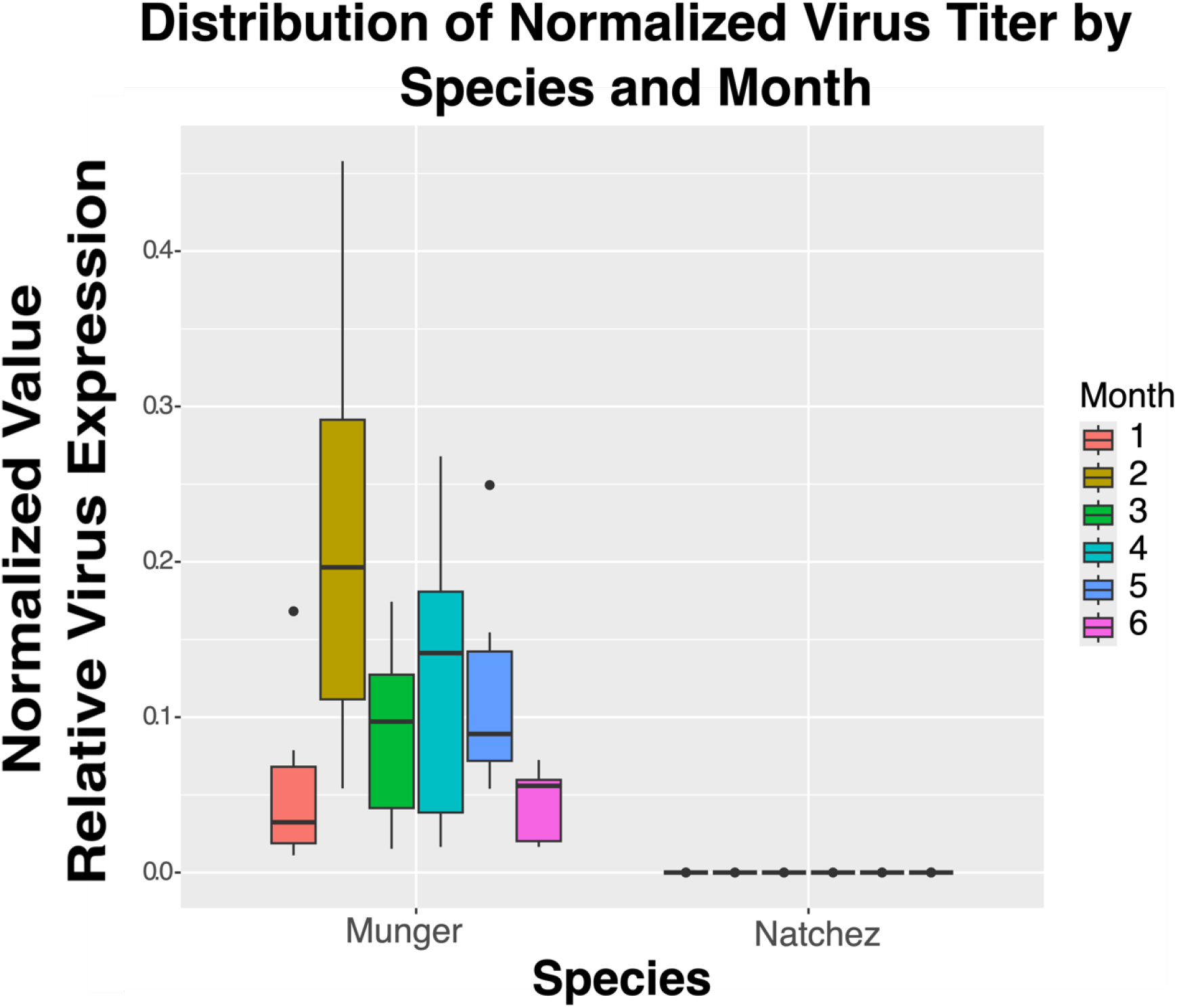
Box plot illustrating the distribution of normalized virus RNA accumulation for blackberry chlorotic ringspot virus (BCRV) in ‘Munger’ black raspberry and ‘Natchez’ blackberry, across a six-month period. Dots represent outliers. Compact letter display (CLD) results showing pairwise comparisons of species and month combinations, where groups sharing the same letter are not significantly different, and groups with different letters are significantly different based on Tukey’s post-hoc test at the 5% level of significance.

## 4. DISCUSSION

This study advances our understanding of BCRV-host interactions by providing insights into the dynamics of virus accumulation over time in two distinct genotypes, ‘Munger’ and ‘Natchez’. We monitored BCRV accumulation over six months, coinciding with the growing season, revealing that ‘Munger’ is favorable host of the virus, consistently supporting higher virus accumulation across all sampling points. In contrast, ‘Natchez’ did not support high virus accumulation, and detection was sporadic throughout the study. These results align with the fact that blackberries are generally more resistant to viral infections than black raspberries (Finn & Clark, 2012). Moreover, these results shed light on why ‘Munger’ has been the standard indicator for virus detection for *Rubus* species.

Ilarviruses typically display uneven distribution and fluctuating titers within woody hosts throughout the year (Uyemoto & Scott, 1992; Zotto et al., 1999). This pattern was evident in ‘Natchez,’ but not in ‘Munger,’ where viral titer levels remained high, facilitating consistent detection. The intermittent detection of the virus in ‘Natchez’ corroborates the observations of Villamor et al (2022), where the authors noted that viruses in berry crops were not consistently detected at four sampling points. The authors hypothesized that this inconsistency could be due to temporal variations in virus accumulation and distribution within the host, as well as the influence of individual host genotypes on virus-host interactions (Villamor et al., 2022). Similar situations have been observed in other plant models, such as sugarcane, where virus detection varies with season and tissue type (Bolus et al., 2023; Malapi-Wight et al., 2021). In our study, we minimized the influence of seasonality and tissue-related titer variation by maintaining consistent temperature and light conditions and sampling newly expanded leaves from both hosts.

Despite the high virus accumulation in ‘Munger’ that was at least 47,300 times greater than in ‘Natchez’, the former remained asymptomatic throughout the study. This situation has also been documented with other viruses infecting *Rubus*, such as blackberry yellow vein associated virus and blackberry virus Y, where ‘Munger’ serves as a host without exhibiting symptoms (Susaimuthu et al., 2007; Martin et al., 2013). Ilarviruses, including BCRV, are known to cause latent infections. While BCRV remains asymptomatic during single infections in blackberry (Poudel et al., 2014) and raspberry, it can transition to a symptomatic state in mixed infections, such as those causing blackberry yellow vein disease.

Such asymptomatic conditions are often associated with latent infections, where a host plant remains symptomless despite a virus infection, potentially due to virus tolerance or viral persistence. Virus tolerance is observed when high viral titers do not trigger a strong antiviral response. In contrast, virus persistence occurs when the virus continues to replicate at low levels over time without inducing disease symptoms (Takahashi et al., 2019). We hypothesize that ‘Munger’ exhibits tolerance to BCRV infection, as it shows no symptoms despite high virus accumulation.

A mixed-effects statistical analysis model was employed to account for repeated measures collected over different months and the variability in observation numbers across groups. This approach effectively addressed the unbalanced data, allowing us to assess the effects of species, month, and their interactions on virus titer accumulation while considering the random effects associated with each individual biological sample. The two-way ANOVA results indicated significant influences from these factors on virus accumulation. Post-hoc analysis revealed higher virus accumulation in ‘Munger’, with viral fluctuating titers across the months and a significant increase during the second sampling point. In contrast, ‘Natchez’ exhibited lower accumulation overall with less variability. The mechanisms underlying these observations remain to be explored.

Plants were subjected to low temperatures for a month before the start of this experiment. Virus replication is believed to slow in cooler conditions, with rising temperatures correlating with increased viral titers, a trend noted in studies linking seasonal changes to viral dynamics (Honjo et al., 2020). This increase is attributed to early viral accumulation in young leaves, with the host’s defense system activating RNA silencing genes as the viral load increases (Honjo et al., 2020). This could explain the significant increase in virus accumulation observed in ‘Munger’ during the second month, likely driven by the temperature shift from 4°C to 22°C. As viral titers peaked, an antiviral response may have been triggered, leading to a decline in virus expression in the following months.

Host genotype significantly influences virus expression, as shown by Smith and Campbell (2004), who studied how time and plant genotype affect the spread of poplar mosaic virus. They found that the virus’ systemic spread varies by genotype, with different *Populus* genotypes exhibiting distinct levels of susceptibility and virus accumulation.

Despite having fewer observations for ‘Munger’ than for ‘Natchez,’ statistical analysis delivered a high level of confidence for several reasons: 1) BCRV was consistently detected at every sampling point in ‘Munger’ unlike ‘Natchez’; 2) ‘Munger’ showed consistent and homogeneous observations, with CT values fluctuating by less than five cycles within each plant (Table 1); and 3) our statistical model accounted for unbalanced data.

Future research should focus on uncovering the host factors that either inhibit or facilitate BCRV’s systemic spread, potentially enhancing our understanding of resistance traits among different *Rubus* genotypes and facilitating their incorporation into breeding programs.

This study offers a detailed analysis of the interaction between blackberry chlorotic ringspot virus (BCRV) and two *Rubus* genotypes, ‘Munger’ and ‘Natchez’, highlighting key differences in virus expression and detection. We found that ‘Munger’ serves as a more permissive host, exhibiting higher and more consistent virus accumulation, making it a valuable indicator for monitoring viruses in *Rubus*. Although both genotypes remained asymptomatic, differences in virus accumulation suggest that further research into host mechanisms could deepen our understanding of virus tolerance and persistence.

## DECLARATIONS

During the preparation of this work the authors used ChatGPT v.4 to assist in generating R code for statistical analysis and to improve the language. No content was directly generated by the tool; rather, it was used to help enhance the overall clarity of the text. After using this tool, the authors reviewed and edited the content and code as needed and take full responsibility for the content of the publication.

## DATA AVAILABILITY STATEMENT

The data supporting the findings of this study are available from the corresponding author.

## ACKNOWLEDGEMENTS

We thank the statistical analysis experts, Dr. Richard Adams and Jenniffer Roa Lozano (University of Arkansas) for helpful discussion on the statistical analysis methodology. IET was supported by the United States National Institute of Food and Agriculture project ARK02850 and the Arkansas Agricultural Experimental Station.

---

**Supplementary Table 1.**
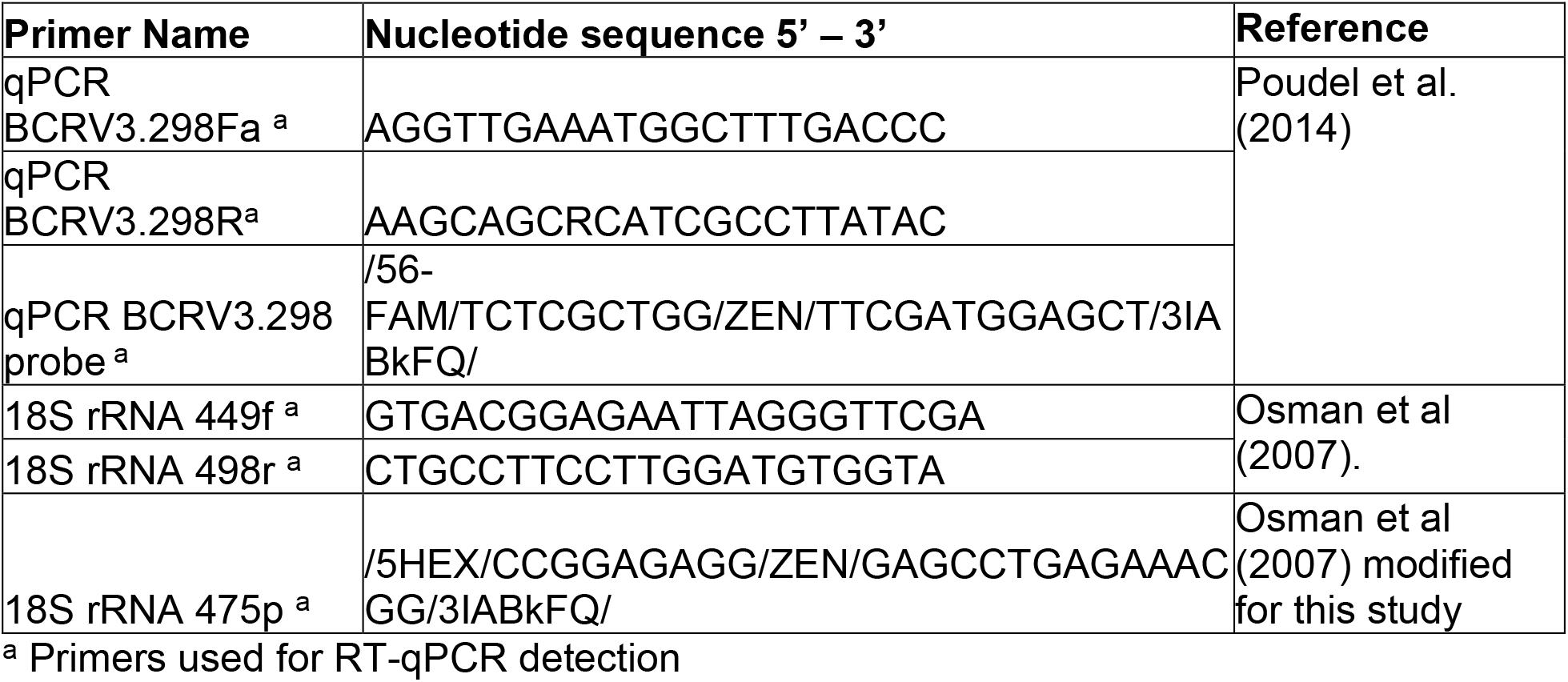
Detection primers and TaqMan™ probes used in this study.

## Supplementary Material

R Code Utilized to perform the statistical analysis in R studio. This code was generated with the aid of ChatGPT v.4

~~~
# Load necessary libraries
install.packages(‘emmeans’)
install.packages(‘lmerTest’)
install.packages(‘multcompView’)
install.packages(‘multcomp’)
install.packages(“pbkrtest”)
library(pbkrtest)
library(lme4) # For mixed-effects models
library(lmerTest) # For p-values in mixed-effects models
library(emmeans) # For post-hoc comparisons
library(ggplot2) # For plotting
library(multcompView)
library(multcomp)
# Load your dataset
# Set working directory and load your dataset
setwd(‘∼/Desktop/’) # Change this to your actual path
# Replace ‘your_data.csv’ with the actual path to your CSV file
data <- read.csv(“Viraltiter6months9.csv”)
# Normalize the CT values, ensuring NA handling
#data$Normalized_Value <- 2 ^ (data$CT_Internal_Control - data$CT_Virus)
# Ensure proper factor levels for species and month
data$Species <- as.factor(data$Species)
data$Month <- as.factor(data$Month)
data$Plant_ID <- as.factor(data$Plant_ID)
# Fit a mixed-effects model, ignoring NA values in the analysis
##model <- lmer(Normalized_Value ∼ Species * Month + (1 | Plant_ID), data = data)
model <- lmer(Normalized_Value ∼ Species * Month + (1 | Plant_ID), data = data)
# Summarize the model
summary(model)
# Check for significant differences using ANOVA
anova(model)
# Post-hoc comparisons
emmeans_results <- emmeans(model, ∼ Species * Month)
#emmeans_results <- emmeans(model, ∼ Species * Month, mode = “asymptotic”)
summary(emmeans_results)
post_hoc_results <- pairs(emmeans_results, adjust = “tukey”)
summary(post_hoc_results)
plot(emmeans_results)
plot(post_hoc_results)
# Convert results to a data frame
post_hoc_df <- as.data.frame(post_hoc_results)
multcomp::cld(emmeans_results)
letters_df <- multcomp::cld(emmeans_results, Letters = letters)
# Get compact letter display
letters_df <- multcomp::cld(emmeans_results, adjust = “sidak”, Letters = letters)
# Compute mean normalized values for each species
mean_munger <- mean(data$Normalized_Value[data$Species == “Munger”], na.rm = TRUE)
mean_natchez <- mean(data$Normalized_Value[data$Species == “Natchez”], na.rm = TRUE)
# Calculate how many times Munger is larger than Natchez
ratio <- mean_munger / mean_natchez
# Print the result
ratio
# Get the unique months from the data
unique_months <- unique(data$Month)
# Initialize a result dataframe to store the ratios for each month
results <- data.frame(Month = character(), Ratio = numeric(),
      Mean_Munger = numeric(), Mean_Natchez = numeric(), stringsAsFactors =
FALSE)
# Loop through each month and calculate the means and ratio
for (month in unique_months) {
 # Compute mean normalized values for Munger in this month
 mean_munger <- mean(data$Normalized_Value[data$Species == “Munger” & data$Month ==
month], na.rm = TRUE)
 # Compute mean normalized values for Natchez in this month
 mean_natchez <- mean(data$Normalized_Value[data$Species == “Natchez” & data$Month ==
month], na.rm = TRUE)
 # Calculate the ratio (how many times Munger is larger than Natchez)
 ratio <- ifelse(mean_natchez == 0, NA, mean_munger / mean_natchez) # Handle division by
zero
 # Append results to the dataframe
 results <- rbind(results, data.frame(Month = month, Ratio = ratio,
            Mean_Munger = mean_munger, Mean_Natchez = mean_natchez))
}
# View the results
print(results)
# Visualization of normalized values over time, handling unbalanced data
ggplot(data, aes(x = Month, y = Normalized_Value, color = Species)) +
 geom_point(position = position_jitter(width = 0.1), alpha = 0.7) + # Add jitter to points for
visibility
 geom_line(aes(group = interaction(Species, Plant_ID)), alpha = 0.5) + # Connect points for
the same plant
 labs(title = “Normalized Viral Load Over Time”, x = “Month”, y = “Normalized Viral Load”) +
 theme_minimal()
# Violin plot of normalized viral load by Species and Month
ggplot(data, aes(x = Month, y = Normalized_Value, fill = Species)) +
 geom_violin(trim = FALSE, alpha = 0.7) + # Violin plot
 geom_boxplot(width = 0.1, position = position_dodge(0.9), outlier.shape = NA, alpha = 0.5) +
# Boxplot on top
 labs(title = “Violin Plot of Normalized Viral Load by Species and Month”,
    x = “Month”,
    y = “Normalized Viral Load”) +
 theme_minimal() +
 theme(legend.title = element_blank())
# Bar Plot
ggplot(data, aes(x = Species, y = Normalized_Value, fill = Month)) +
 geom_bar(stat = “summary”, fun = “mean”, position = “dodge”) +
 labs(title = “Normalized Virus Titer by Species and Month”, y = “Normalized Value”)
# Box Plot
ggplot(data, aes(x = Species, y = Normalized_Value, fill = Month)) +
 geom_boxplot() +
 labs(title = “Distribution of Normalized Virus Titer by Species and Month”, y = “Normalized
Value”)
~~~

## REFERENCES

Ammara, U., Al-Sadi, A. M., Al-Shihi, A., & Amin, I. (2017). Real-time qPCR assay for the TYLCV titer in relation to symptoms-based disease severity scales. Int. J. Agric. Biol, 19, 145–151. 10.17957/IJAB/15.0256

Bin, Y., Xu, J., Duan, Y., Ma, Z., Zhang, Q., Wang, C., … & Zhou, C. (2022). The titer of citrus yellow vein clearing virus is positively associated with the severity of symptoms in infected citrus seedlings. Plant Disease, 106(3), 828–834. 10.1094/PDIS-02-21-0232-RE

Bolus, S., Wathen-Dunn, K., Grinstead, S. C., Hu, X., Malapi, M., & Mollov, D. (2023). Effects of tissue type and season on the detection of regulated sugarcane viruses by high throughput sequencing. CABI Agriculture and Bioscience, 4(1), 34. 10.1186/s43170-023-00175-1

Druciarek, T., Sierra-Mejia, A., Zagrodzki, S. K., Singh, S., Ho, T., Lewandowski, M., & Tzanetakis, I. E. (2024). Phyllocoptes parviflori is a distinct species and a vector of the pervasive blackberry leaf mottle associated virus. Infection, Genetics and Evolution, 117, 105538. 10.1016/j.meegid.2023.105538

Finn, C.E. & Clark, J.R. (2012). Blackberry. In M. Badenes & D. Byrne (Eds.), Fruit breeding (Vol. 8, pp. 151–190). Springer. 10.1007/978-1-4419-0763-9_5

Hedrick, U. P. (1925). The small fruits of New York. Report of the New York State Agricultural Experiment Station for the year ending June 30, 1925, pt. II. J.B. Lyon Company.

Honjo, M. N., Emura, N., Kawagoe, T., Sugisaka, J., Kamitani, M., Nagano, A. J., & Kudoh, H. (2020). Seasonality of interactions between a plant virus and its host during persistent infection in a natural environment. The ISME journal, 14(2), 506–518. 10.1038/s41396-019-0519-4

Malapi-Wight, M., Adhikari, B., Zhou, J., Hendrickson, L., Maroon-Lango, C. J., McFarland, C., Foster, J.A., & Hurtado-Gonzales, O. P. (2021). HTS-based diagnostics of sugarcane viruses: Seasonal variation and its implications for accurate detection. Viruses, 13(8), 1627. 10.3390/v13081627

Martin R.R., Constable F., & Tzanetakis I.E. (2016) Quarantine Regulations and the Impact of Modern Detection Methods. Annual Review Phytopathology. 54, 189–205. 10.1146/annurev-phyto-080615-100105.

Martin, R.R. & Tzanetakis, I.E. (2008). Characterization of three novel viruses infecting raspberry. Acta Horticulturae. 777, 317–322 10.17660/ActaHortic.2008.777.47

Martin, R. R., MacFarlane, S., Sabanadzovic, S., Quito, D., Poudel, B., & Tzanetakis, I. E. (2013). Viruses and virus diseases of Rubus. Plant disease, 97(2), 168–182. 10.1094/PDIS-04-12-0362-FE

Osman, F., Leutenegger, C., Golino, D., & Rowhani, A. (2007). Real-time RT-PCR (TaqMan®) assays for the detection of Grapevine Leafroll associated viruses 1–5 and 9. Journal of virological methods, 141(1), 22–29. 10.1016/j.jviromet.2006.11.035

Poudel, B. (2011). Epidemiological studies on Blackberry yellow vein associated virus and Blackberry chlorotic ringspot virus. M.Sc. thesis. University of Arkansas, Fayetteville.

Poudel, B., Ho, T., Laney, A., Khadgi, A., & Tzanetakis, I. E. (2014). Epidemiology of Blackberry chlorotic ringspot virus. Plant disease, 98(4), 547–550. 10.1094/PDIS-08-13-0866-RE

Quito-Avila, D. F., & Martin, R. R. (2012). Real-time RT-PCR for detection of Raspberry bushy dwarf virus, Raspberry leaf mottle virus and characterizing synergistic interactions in mixed infections. Journal of virological methods, 179(1), 38–44. 10.1016/j.jviromet.2011.09.016

Sierra-Mejia A. (2024). Advancing molecular tools for the study of Rubus viruses. PhD dissertation submitted at the University of Arkansas, pp.118

Sierra-Mejia A., Villamor D.E.V., & Tzanetakis I.E. (2025). Development and application of an infectious clone and gene silencing vector derived from blackberry chlorotic ringspot virus. Virus Research. 10.1016/j.virusres.2024.199460

Smith, C. M., & Campbell, M. M. (2004). Populus genotypes differ in infection by, and systemic spread of, Poplar mosaic virus. Plant pathology, 53(6), 780–787. 10.1111/j.1365-3059.2004.01095.x

Susaimuthu, J., Gergerich, R. C., Bray, M. M., Clay, K. A., Clark, J. R., Tzanetakis, I. E., & Martin, R. R. (2007). Incidence and ecology of Blackberry yellow vein associated virus. Plant Disease, 91(7), 809–813. 10.1094/PDIS-91-7-0809

Takahashi, H., Fukuhara, T., Kitazawa, H., & Kormelink, R. (2019). Virus latency and the impact on plants. Frontiers in Microbiology, 10, 2764. 10.3389/fmicb.2019.02764

R Core Team. (2023). R: A language and environment for statistical computing (Version 4.4.1). R Foundation for Statistical Computing. https://www.R-project.org/

Uyemoto, J. K., & Scott, S. W. (1992). Important diseases of Prunus caused by viruses and other graft-transmissible pathogens in California and South Carolina. Plant Disease, 76(1), 5–11.

Villamor, D. E. V., Keller, K. E., Martin, R. R., & Tzanetakis, I. E. (2022). Comparison of high throughput sequencing to standard protocols for virus detection in berry crops. Plant disease, 106(2), 518–525. 10.1094/PDIS-05-21-0949-RE

Zotto, A. D., Nome, S. F., Di Rienzo, J. A., & Docampo, D. M. (1999). Fluctuations of Prunus necrotic ringspot virus (PNRSV) at various phenological stages in peach cultivars. Plant Disease, 83(11), 1055–1057.

